# Slow to Start, Free at Last: Dual Effects of Mucin on *Escherichia coli* Phage T4

**DOI:** 10.64898/2026.05.21.726763

**Authors:** Kira Céline Koonce, Anna Rasmussen, Raquel Bastos Gonçalo, Mathias Middelboe, Karina B. Xavier, Jesper Juel Mauritzen, Nina Molin Høyland-Kroghsbo

**Affiliations:** University of Copenhagen, Department of Plant and Environmental Sciences, Frederiksberg, Denmark; Gulbenkian Institute for Molecular Medicine, Oeiras, Portugal; University of Copenhagen, Department of Biology, Helsingør, Denmark; Faculdade de Medicina da Universidade de Lisboa, Lisbon, Portugal

## Abstract

Bacteriophages traversing the gastrointestinal tract are exposed to extreme physicochemical stresses that may rapidly compromise virion integrity and shape infection dynamics. While some phages bind to host-derived mucins at mucosal surfaces, the functional consequences of soluble mucin glycans for phage-host interactions remain incompletely understood. Here, we show that soluble mucin glycans exert dual effects on the *Escherichia coli* phage T4 by delaying infection initiation while simultaneously providing environmental virion stability. Mucin-coated T4 exhibits a lag in the onset of productive infection, consistent with transient steric occlusion from *E. coli*, yet without impairing overall phage progeny production once infection was established. We further show that *E. coli* can metabolize purified mucin, supporting a model in which dynamic remodeling of the mucin matrix gradually releases T4 and enables infection. Importantly, mucin coating substantially increases T4 survival under gastrointestinal-like stresses, including acidic pH and protease exposure. Moreover, we find that in a murine gut colonization model, a single oral dose of mucin-coated T4 displayed enhanced fecal persistence over a two-week period, which correlated with prolonged suppression of *E. coli* populations and delayed resolution of phage-associated functional shifts in the gut microbiome. Together, we find that that soluble mucin glycans actively shape T4 phage infection kinetics, virion stability, and ecological impact in the murine gut, and support mucin-based formulations as a strategy to extend the persistence and efficacy of orally delivered phages.

## Introduction

Bacteriophages (phages) are the most abundant biological entities on Earth and are widespread across microbial ecosystems, including those associated with the human body [1]. As natural predators of bacteria, they shape microbial community structure, population dynamics, and evolutionary trajectories, and are increasingly recognized to participate in additional interactions within eukaryotic host-associated environments beyond their bacterial targets [2].

Phage lifecycles fall broadly into lytic and temperate modes. Lytic phages infect and ultimately kill their hosts through intracellular replication followed by cell lysis, whereas temperate phages integrate into the bacterial chromosome as prophages, through which they modulate host physiology and contribute to horizontal gene transfer [3]. Among lytic phages, *Escherichia coli* phage T4 represents a well-studied model for virulent infection. T4 initiates infection by its tail fibers reversible binding to the *E. coli* outer membrane porin OmpC and lipopolysaccharide [4]. This triggers conformational changes, leading to irreversible attachment, and eventually to phage genome injection. Generally, bacteria and phages engage in a continual evolutionary arms race [5]. Bacteria deploy multilayered defense systems, including receptor modification or receptor loss, CRISPR-Cas immunity, and abortive infection pathways, to limit or block phage infection [5]. In response, phages evolve counterstrategies such as diversification of receptor-binding proteins, anti-CRISPR factors, and other mechanisms that suppress bacterial defense.

While phage-bacterial interactions are often characterized under controlled laboratory conditions, encounters *in vivo* unfold within far more complex, structured environments that impose spatial constraints, biochemical stressors, and involve tripartite (phage-bacterial-eukaryotic host) interactions capable of substantially altering outcomes observed in simplified systems [6]. In the gastrointestinal (GI) tract, such interactions unfold in the context of the gut microbiota, a dense and dynamic community of microorganisms that colonize the GI tract in a mutually beneficial relationship with the host [7, 8]. The composition and diversity of this community are shaped by host lifestyle, diet, morphology, phylogeny, and long-term host-microbe co-evolution [9], and the microbiota plays essential roles in host metabolism, pathogen defense, and nervous system function. Within this complex ecosystem, diverse bacterial populations co-exist alongside abundant phages, creating a setting in which phage-bacterial interactions are frequent, dynamic, and highly context dependent [10, 11].

One of the key factors shaping the microbiota function at epithelial surfaces is mucus. Phages encounter mucus across multiple epithelial systems, including the respiratory, urogenital, and GI tracts [12]. Among these, the GI mucosa represents a particularly dynamic setting, rich in phage-bacterial encounters. Mucin, the principal polymeric component of mucus, is a heavily O-glycosylated, gel-forming glycoprotein that can self-assemble into viscoelastic networks through self-aggregation, disulfide crosslinking, and noncovalent interactions [13]. In the GI tract, both secreted and membrane-tethered mucins form a barrier that structures microbial habitats, regulates water and solute transport, and helps maintain a balanced microenvironment at the epithelial surface, for example, by limiting direct exposure to extreme pH. These selective transport and retention properties of mucin gels shape how microorganisms navigate the mucosal interface. Importantly, this mucus layer is characterized by continuous secretion, turnover, and enzymatic cleavage, remodeling the gel and releasing mucin-derived O-glycans and fragments into the surrounding fluid [14]. These soluble, free-floating mucin glycans are abundant in the gut, where they are metabolized by specific commensal bacteria and contribute to community structure.

In light of these complexities, the Bacteriophage Adherence to Mucus (BAM) model illustrates how mucus-producing epithelial surfaces influence phage ecology by shaping phage behavior in ways that benefit both phages - through increased encounters with bacterial hosts - and the eukaryotic host - through enhanced mucosal phage-mediated pathogen defense [15]. Within this framework, surface-associated mucins are not passive barriers but actively shape how phages localize and move within mucus layers at epithelial interfaces. Certain phages carry capsid-associated Ig-like domains that mediate reversible binding to mucin glycans; for example, the T4 outer capsid protein Hoc binds mucin-associated glycans, resulting in phage accumulation within epithelial mucus layers and constrain motion in this environment [16].

Building on the BAM model, mucins distinctly affect phage-host interactions in different phage-host systems. In particular, mucins have been shown to modulate bacterial physiology, virulence, susceptibility to phage infection, and phage resistance development across multiple systems, including *Flavobacterium columnare* [17, 18], *Streptococcus mutans* [19] and *Yersinia enterocolitica* [20]. Further, mucin-phage interactions have been associated with enhanced short-term phage persistence accompanied by antimicrobial effects in mucosal environments *in vitro* and in mice [21].

While phage adherence to mucus is well established as a mechanism that increases phage-bacterial encounter rates at mucosal surfaces, it remains unclear how mucin interactions influence early steps of infection, such as the timing of phage adsorption to bacterial hosts, independent of their effects on overall infection outcome. At the same time, mucins are well established as central protective components of mucosal barriers, and phages are known to interact directly with mucin glycans [13, 15]. However, evidence for mucin-mediated protection of phages has so far largely been inferred from increased binding within mucus layers, rather than from direct effects on phage stability. To date, the extent to which mucins directly affect phage stability over prolonged time under physiologically relevant GI stresses remains underexplored. Here, we investigate how mucin coating of T4 modulates T4-*E. coli* interactions, focusing on early infection dynamics and phage stability under *in vitro* GI-like stress coupled with determining T4 persistence over a two-week period, as well as *E. coli* killing, and overall microbiome effects *in vivo* in the murine gut.

## Results

### Mucin coating delays T4 adsorption without impairing replication capacity

While T4 phages can accumulate within mucus layers through reversible interactions between capsid-associated Hoc proteins and mucin glycans [15], it remains unclear how mucin influences the kinetics of phage-host interactions once phages encounter bacteria. To assess early infection kinetics, we treated *E. coli* MG1655 with control-treated or T4 pre-coated in 1% mucin type II and quantified free T4 phages over time by enumerating plaque-forming units, PFU/mL (**Fig. 1A**). Under control conditions, phage abundance rose 10-fold at 30 min, consistent with the established 25-30 min latent period of T4 [22]. At 90 min, replication peaked, visible as the highest fold increase in free phage within each 30-min interval (**Fig. 1B**), before reaching a plateau by 120 min. In contrast, when coated in mucin, free T4 levels remained steady until 60 min, with peak replication shifted to 120 min and a plateau reached around 150 min. Despite this shift, the height of the replication peak was comparable to the control, indicating that mucin does not impair replication or lysis once infection begins, but primarily delays the initiation of productive infection.

**Figure 1.**
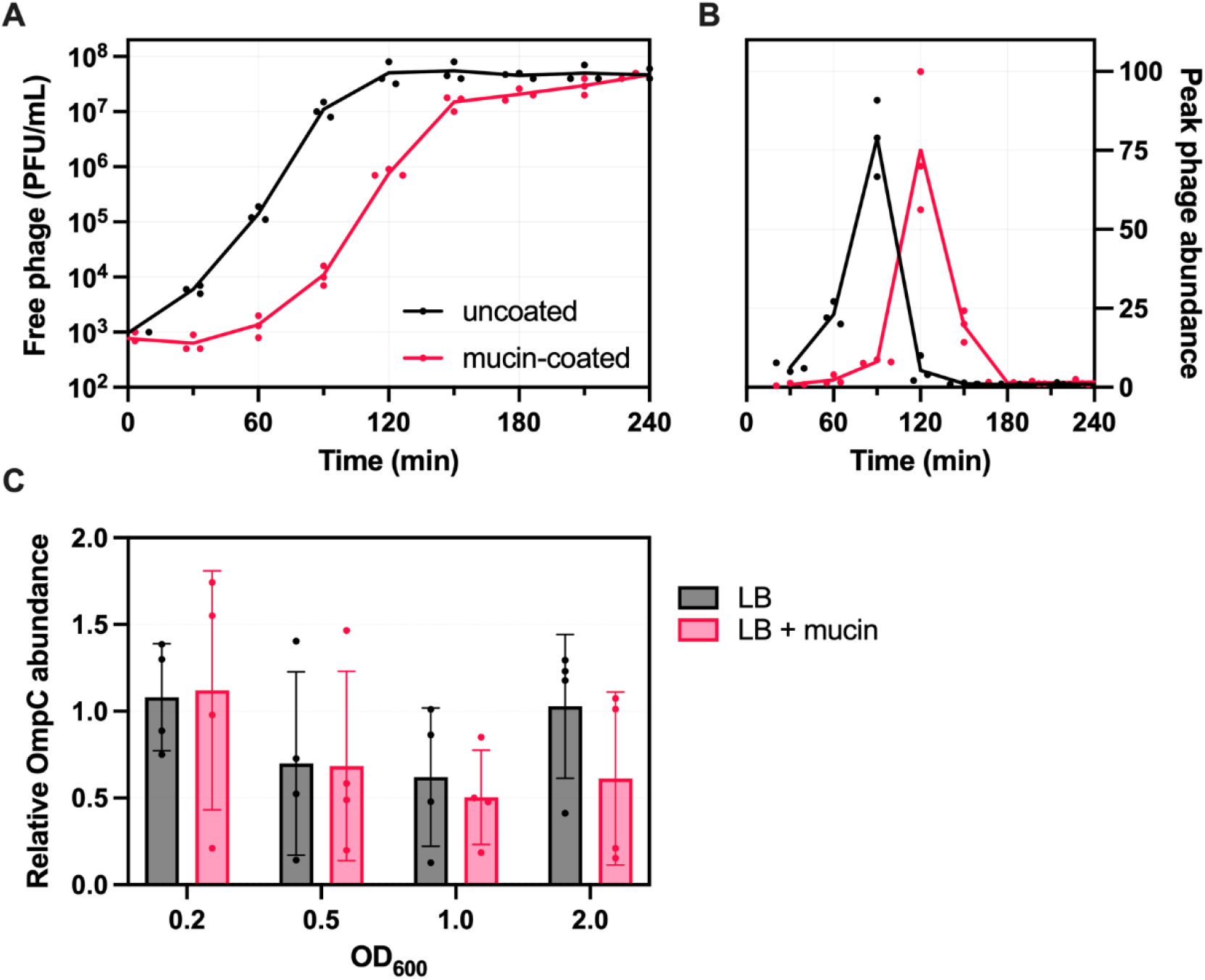
Mucin delays phage-bacterial infection dynamics without impairing phage replication. To assess the effect of mucin-coating on early T4 infection dynamics, we exposed *E. coli* MG1655 to uncoated or mucin-coated T4 and quantified the abundance of free T4 over time. **(A)** Free phage measured over time as PFU/mL following addition of uncoated or mucin-coated T4 to *E. coli* MG1655 cultures. *n* = 3. **(B)** Peak phage abundance, calculated as the fold-increase in free phage over each 30-min interval (PFU_t_/PFU_t-30min_) as a measure of phage replication efficiency. *n* = 3. **(C)** Relative OmpC protein levels in *E. coli* grown in LB or LB supplemented with 1% mucin, measured by Western Blotting and normalized to RpoB at the indicated OD_600_. Error bars represent mean ± SD from *n* = 4 biological replicates.

We considered two mechanistic routes that could explain why mucin delays T4 replication: either mucin alters the physiological state of the *E. coli* host or mucin directly alters the state of the phage, e.g. by temporarily masking phage interaction with the host. To assess whether mucin alters *E. coli* in a way that could influence phage infection dynamics, we quantified the abundance of OmpC, the primary T4 phage receptor [4]. We measured OmpC levels across cell densities by Western blotting and normalized protein abundance to RpoB. OmpC levels were indistinguishable between control- and mucin-treated *E. coli* cultures, demonstrating that mucin exposure did not alter receptor abundance under our experimental conditions (**Fig. 1C**). Thus, altered phage receptor regulation does not account for the delayed mucin-mediated onset of T4 infection.

We next examined whether mucin instead occludes T4 virions, thereby transiently blocking the ability of the phage tail fibers to access the *E. coli* surface receptors required for adsorption. Mucin binds Hoc proteins on the T4 capsid, and the mucin glycopolymers themselves readily self-associate into extended networks in solution [13, 15]. Together, these properties might generate a polymeric coating around the virion that sterically restricts tail-fiber mobility and temporarily prevents productive receptor engagement. Given the temporary nature of the adsorption delay, which indicates that mucin does not impose a permanent physical obstruction, we hypothesized that the mucin coating surrounding phage particles is progressively relieved over time.

We were curious if *E. coli* MG1655 could actively remodel or degrade the mucin coating, thereby progressively releasing trapped T4 and enabling their adsorption. Thus, we tested whether *E. coli* MG1655 can use mucin as sole carbon source. While it did not grow in a carbon-free control M9, it increased in biomass over time in M9 supplemented with mucin, demonstrating its ability to effectively metabolize mucin type II (**Fig. 2A**). To visualize bacterial behavior and spatial mucin organization under these conditions, we performed time-lapse phase-contrast microscopy of *E. coli* MG1655 grown in M9 supplemented with mucin (**Movie S1**). Representative snapshots from the beginning of the experiment and after 10 h of growth reveal *E. coli* MG1655 cells together with heterogeneous mucin aggregates up to ∼18 µm in size that are visible at both time points. Bacterial cell density increased over time, consistent with bacterial survival and growth in mucin-supplemented medium (**Fig. 2B**). In parallel, we used negative-stain transmission electron microscopy (TEM) to qualitatively assess the spatial relationship between T4 phages and mucin. Phage capsids were observed embedded in dense, filamentous structures consistent with mucin-derived aggregates (**Fig. S1**). These images provide visual support for a spatial association between virions and the mucin matrix, although negative-stain TEM visualizes overall mass distribution rather than native structure or direct molecular interactions.

**Figure 2.**
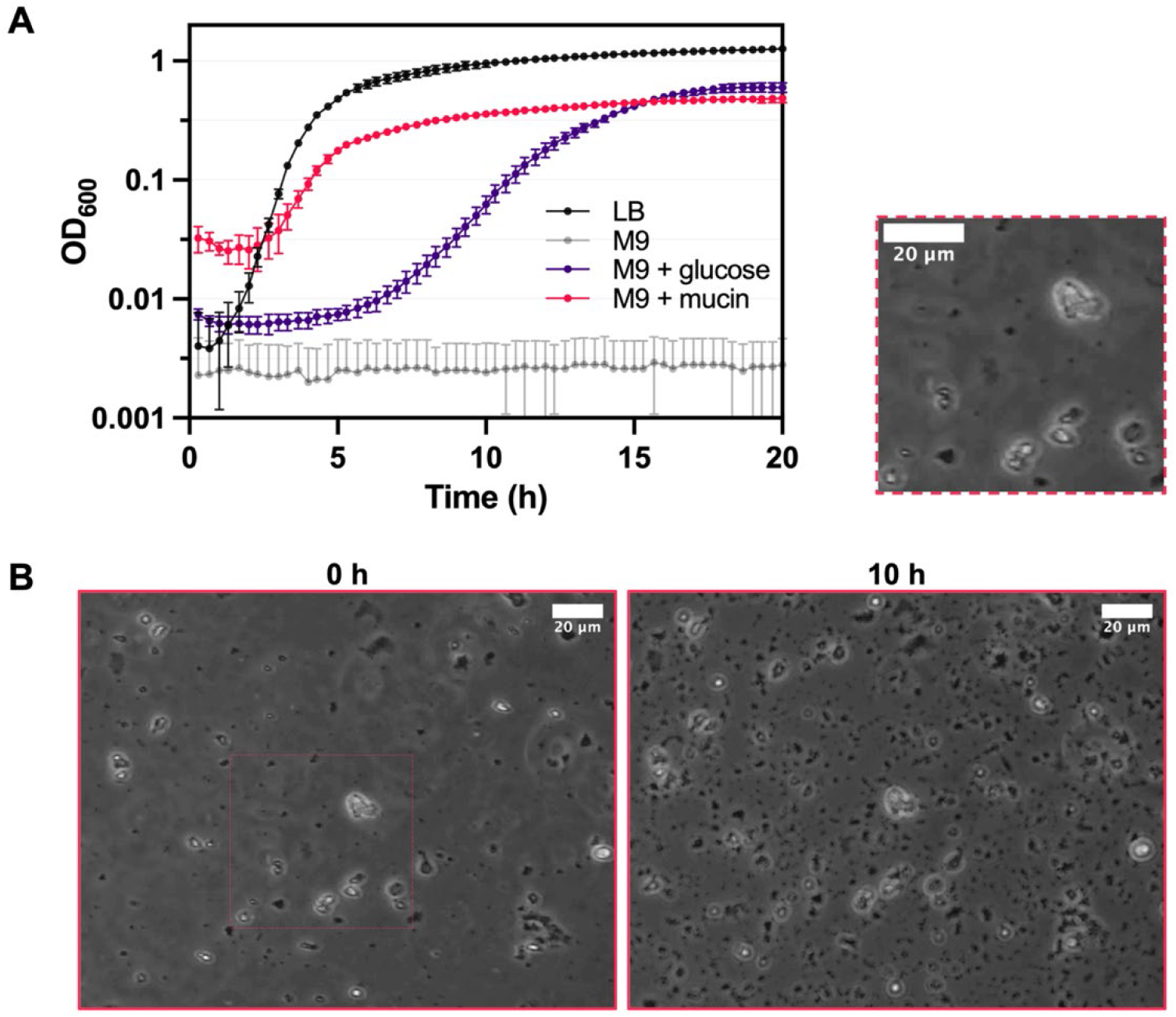
*E. coli* MG1655 metabolizes mucin type II as sole carbon source. **(A)** To assess the ability of *E. coli* MG1655 to actively alter mucin, cells were either grown in LB as control or subjected to different M9 minimal medium treatments with or without carbon supplementation of 0.4% glucose or 1% mucin and grown for 20 h at 37 °C. The OD_600_ is displayed after subtracting the respective solvent blanks. Error bars represent mean ± SD from *n* = 3 biological replicates. **(B)** Representative phase-contrast microscopy images of *E. coli* MG1655 grown in M9 medium supplemented with 1% mucin at 0 h and after 10 h of incubation at room temperature. Mucin aggregates are visible as ∼1-18 µm irregularly structures. An enlarged subpanel highlights a selected region of the 0 h image (as indicated). Images correspond to selected frames from the time-lapse experiment shown in Movie S1. Magnification: 40×, scale bar: 20 µm.

Together, these observations support a model in which T4 phages are transiently occluded within the mucin matrix, limiting initial adsorption to *E. coli*. Over time, this constraint is relieved, allowing virions to access bacterial receptors and initiate productive infection. The ability of *E. coli* MG1655 to metabolize mucin is consistent with bacterial processing as a plausible contributor to remodeling of the mucin matrix over time, which may facilitate relief of phage occlusion. This occlusion-and-release mechanism may explain the delayed onset of T4 replication while remaining consistent with the unchanged burst magnitude once infection is established.

### Mucin coating enhances T4 stability under gastrointestinal-like stresses

Having established that mucin coating delays the onset of productive infection without impairing replication output (**Fig. 1**), we next asked whether the protective role of mucus extends beyond spatial localization effects described in the BAM model to directly influence the phage particle itself [15]. However, whether mucins also confer direct protection of T4 phages against physicochemical stress has remained largely unexplored. In the GI tract, mucus forms the first barrier against environmental stress, shielding both microbes and host tissue from acidic pH, digestive enzymes, and other destabilizing conditions [13]. Because T4 naturally binds mucin glycopolymers, we hypothesized that this interaction might similarly buffer virions against GI-like stress.

To test this, we compared the stability of uncoated and mucin-coated T4 exposed to pH 3 and to the digestive protease α-chymotrypsin (**Fig. 3**), quantifying phage viability as the recovery of infectious particles measured by PFU/mL. Phages were incubated for 1 h at pH 3 to mimic transient gastric passage and for up to 48 h in α-chymotrypsin to model prolonged proteolytic exposure within the intestinal lumen [23]. Initial T4 counts were indistinguishable between uncoated and mucin-coated phage, as expected. Under neutral pH and in the absence of degradative enzymes, uncoated T4 phages remained stable over time, with comparable recoveries to mucin-coated T4 (**Fig. 3A-B**). Under at pH 3, the viability of uncoated T4 dropped to 51% after 30 min and 18% after 60 min (**Fig. 3C**). In contrast, mucin-coated T4 retained 81% viability at 30 min and 52% at 60 min, demonstrating that mucin protects T4 against acidic stress. When exposed to α-chymotrypsin, uncoated T4 phages retained 56% viability after 24 h and 16% after 48 h, whereas mucin-coated T4 phages retained 85% viability at 24 h and 38% at 48 h (**Fig. 3D**), showing that mucin protects T4 against proteolysis.

**Figure 3.**
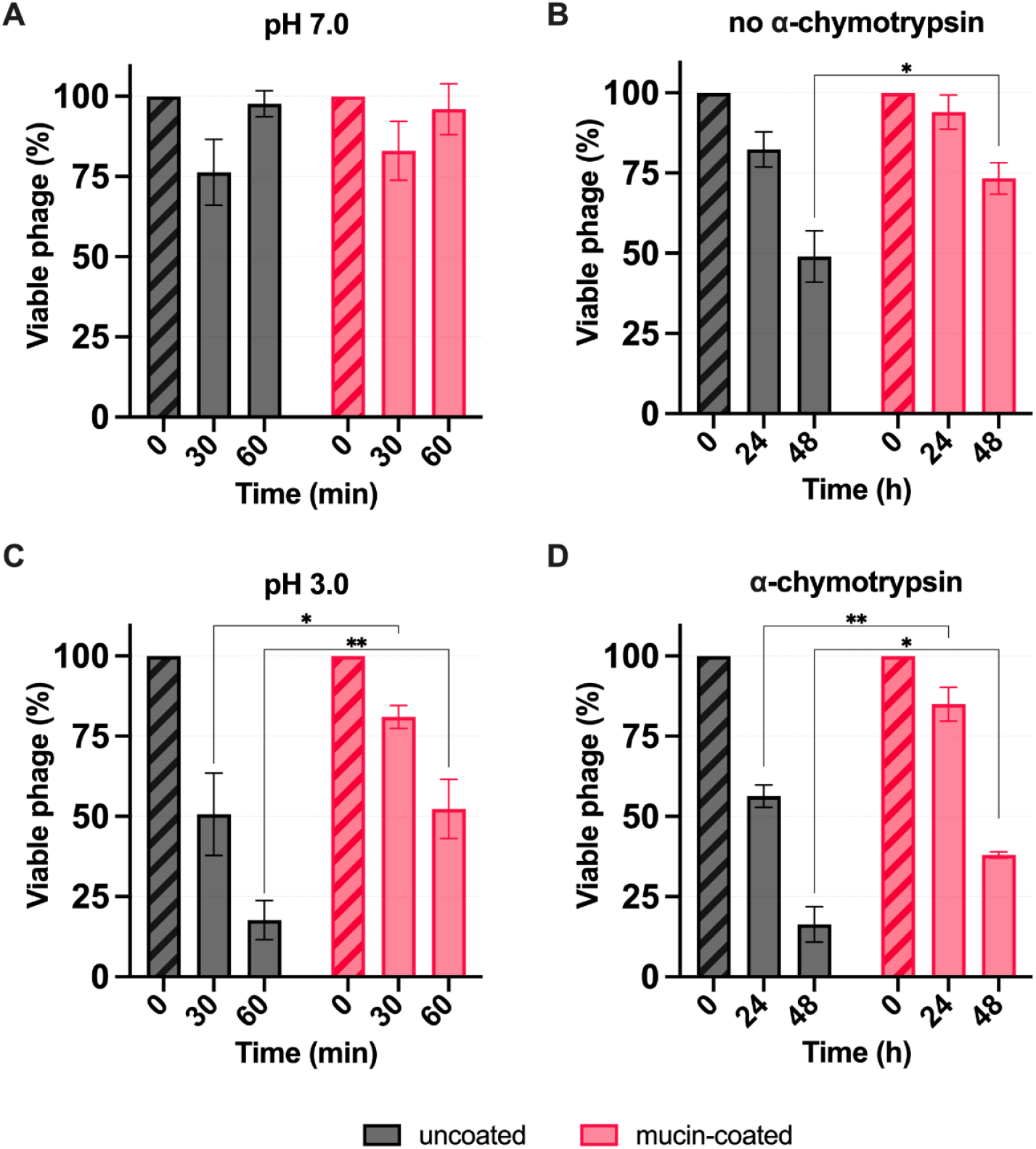
Mucin-coating protects T4 against gastrointestinal-like stressors. To assess whether mucin may protect T4 phages during GI transit, the viability of uncoated and mucin-coated T4 phages was measured relative to *t* = 0 following exposure to GI-relevant stressors *in vitro*. Panels A and B show control conditions. (A) Phages were incubated at neutral pH 7.0 for 30-60 min, and (B) maintained in buffer at 37 °C without α-chymotrypsin for 24-48 h to assess inherent phage stability. Panels C and D model physiologically relevant conditions under gastric and small-intestinal transit. (C) Phages were exposed to acidic conditions at pH 3.0 for 30-60 min to mimic the stomach environment, and (D) incubated with α-chymotrypsin for 24-48 h to simulate proteolytic stress in the small intestine. Error bars represent mean ± SD from *n* = 3 biological replicates. Statistical significance was determined using a mixed-effects model followed by Tukey’s multiple-comparisons test.

These data show that mucin coating provides increased phage stability under both acidic and proteolytic stress. Within the BAM framework, mucin-phage interactions have primarily been linked to phage retention at mucosal surfaces, with protection inferred from spatial localization rather than direct stabilization of the virion [15]. Our findings extend this view by demonstrating that mucin enhances survival of phage particles under GI-like stress conditions. Although α-chymotrypsin is a broad digestive protease that cleaves accessible peptide backbones [23], type II mucin used here is highly glycosylated, with dense O-linked glycans attached to serine and threonine residues. These glycans extend outward from the peptide core and sterically block protease access to the underlying aromatic residues that α-chymotrypsin requires for cleavage. As a result, the mucin backbone is likely effectively shielded from enzymatic attack and remains stable even under prolonged GI-like stresses, allowing the mucin coating to persist as a protective shield for the phage.

### Mucin coating enhances T4 recovery during murine gastrointestinal passage

To determine whether the protective effect of mucin coating observed *in vitro* extends to GI passage *in vivo*, we examined T4 abundance and persistence in a murine gut *E. coli* colonization model (**Fig. 4A**). We treated specific pathogen-free (SPF) mice with streptomycin to permit and maintain stable intestinal colonization by *E. coli* and we subsequently colonized the mice with *E. coli* MG1655 strep^R^. After colonization was established, we gavaged mice with a single oral dose of 100 µL containing 10^7^ PFU uncoated T4, mucin-coated T4 (1 mg total mucin per mouse), or 1x PBS as a control. We collected fecal pellets over a two-week period following gavage and quantified T4 phage abundance.

**Figure 4.**
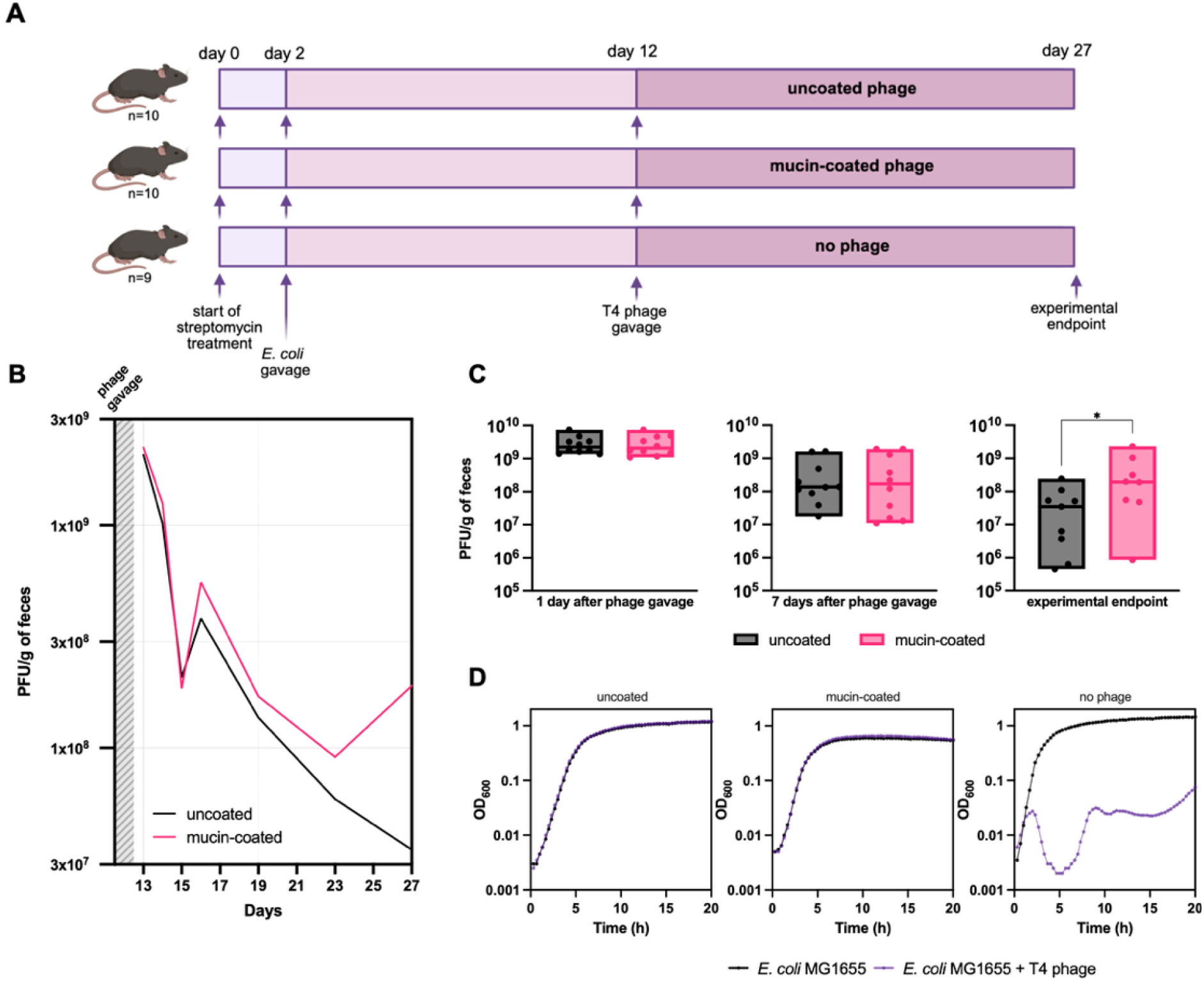
Mucin coating increases T4 persistence in the murine gastrointestinal tract. To assess whether the protective effect of mucin-coating of T4 transfers to GI passage *in vivo*, we orally gavaged uncoated and mucin-coated phages to mice and quantified their abundance and persistence over time. **(A)** Schematic overview of the mouse experiment.Mice were continuously exposed to streptomycin-containing drinking water, pre-colonized with *E. coli* MG1655 on day 2 by oral gavage and subsequently gavaged with either uncoated T4 phage, mucin-coated T4 phage, or PBS, on day 12. Fecal samples were collected at defined timepoints until the experimental endpoint (day 27) to quantify phage persistence over time.**(B)** Fecal phage titers (PFU/g) across the experimental sampling window, displayed as median values for uncoated T4 (*n* = 10) and mucin-coated T4 (*n* = 10) treatment groups. Gray shading indicates the timing of phage gavage. **(C)** Fecal phage titers (PFU/g) shown for individual mice 1 day after phage gavage (day 13), 7 days after (day 19), and on the experimental endpoint (day 27). Horizontal lines indicate the median with interquartile range. Statistical significance was assessed using a Mann-Whitney test. **(D)** Representative growth curves of *E. coli* isolates recovered from fecal samples 1 day after phage treatment (day 13) across uncoated T4, mucin-coated T4, and no phage control treatments, tested for susceptibility to T4 phage.

At day 1 following oral gavage of 100 µL containing a total of 10^7^ PFU uncoated or mucin-coated T4, we recovered approximately 3 × 10^9^ PFU/g feces in both treatment groups (**Fig. 4B-C**), indicating initial substantial phage amplification in the gut regardless of mucin treatment. Importantly, while mucin coating applied only to the initially gavaged phage particles, the majority of phages recovered from feces at later timepoints are expected to be new, uncoated progeny generated during rounds of subsequent replication. Fecal T4 abundance subsequently dropped by over 10-fold between day 1 and day 3 after phage treatment (experimental days 13 and 15) (**Fig. 4B**). Over the following days, we consistently recovered T4 at higher levels in the treatment group initially gavaged with mucin-coated T4, compared to the treatment group receiving uncoated T4 (**Fig. 4B**). At the experimental endpoint 15 days after phage treatment (experimental day 27), we recovered approximately 2 × 10^8^ PFU/g T4 for the treatment group initially receiving mucin-coated T4 compared with 3.5 × 10^7^ PFU/g for the treatment group initially receiving uncoated T4, indicating that pre-coating T4 with mucin before oral administration provides an approximately 6-fold increased phage recovery rate after 15 days (*p* = 0.0464) (**Fig. 4C**).

The higher T4 recovery in the mice receiving mucin-coated T4 may be explained by enhanced phage replication output in the gut, increased phage survival during GI passage, or a combination of both. Because enhanced phage replication would require the continued presence of susceptible hosts, we assessed T4 susceptibility of *E. coli* MG1655 recovered from feces to evaluate whether replication could account for the observed phage recovery. Isolates obtained 1 day after phage gavage (experimental day 13) from both phage-treated groups showed no detectable growth inhibition in the presence of T4, whereas isolates from PBS-treated controls remained phage-susceptible (**Fig. 4D**). Accordingly, mucin-mediated protection of T4, as demonstrated under GI-like stress conditions *in vitro* (**Fig. 3**), likely contributes to the higher levels of recoverable phage observed *in vivo* (**Fig. 4**), yet this factor alone cannot explain the prolonged, increased levels of T4 in pellets from mice treated with mucin-coated T4.

### Mucin-coating prolongs T4-mediated reduction in *E. coli* abundance and modifies the fecal microbiome

We next examined the effect of mucin coating of T4 on *E. coli* colonization and the overall gut microbiome composition. Prior to phage treatment, *E. coli* MG1655 stably colonized the gut at approximately 5 × 10^8^ CFU/g of feces (experimental day 12). One day after phage administration (experimental day 13), fecal *E. coli* abundance was reduced by approximately 4 log_10_ CFU/g in both phage-treated groups relative to control treated animals (*p* = 0.0002 for uncoated T4 and *p* = 0.0022 for mucin-coated T4 treatment; **Fig. 5A**), indicating a strong early T4-mediated *E. coli* killing. Although overall *E. coli* MG1655 levels rose in all groups over time, mice receiving mucin-coated T4 maintained lower *E. coli* burden at later timepoints. 7 days after phage treatment, fecal *E. coli* abundance in the mice receiving uncoated T4 had recovered to similar levels as the control treated mice (*p* = 0.6307). In contrast, *E. coli* levels in the mice that had received mucin-coated T4 remained reduced at day 7 compared to the mice receiving control treatment and treatment with uncoated T4 (*p* = 0.0423; **Fig. 5B**). These differences in *E. coli* dynamics are consistent with the enhanced phage recovery observed in the mice receiving mucin-coated T4 (**Fig. 4**).

**Figure 5.**
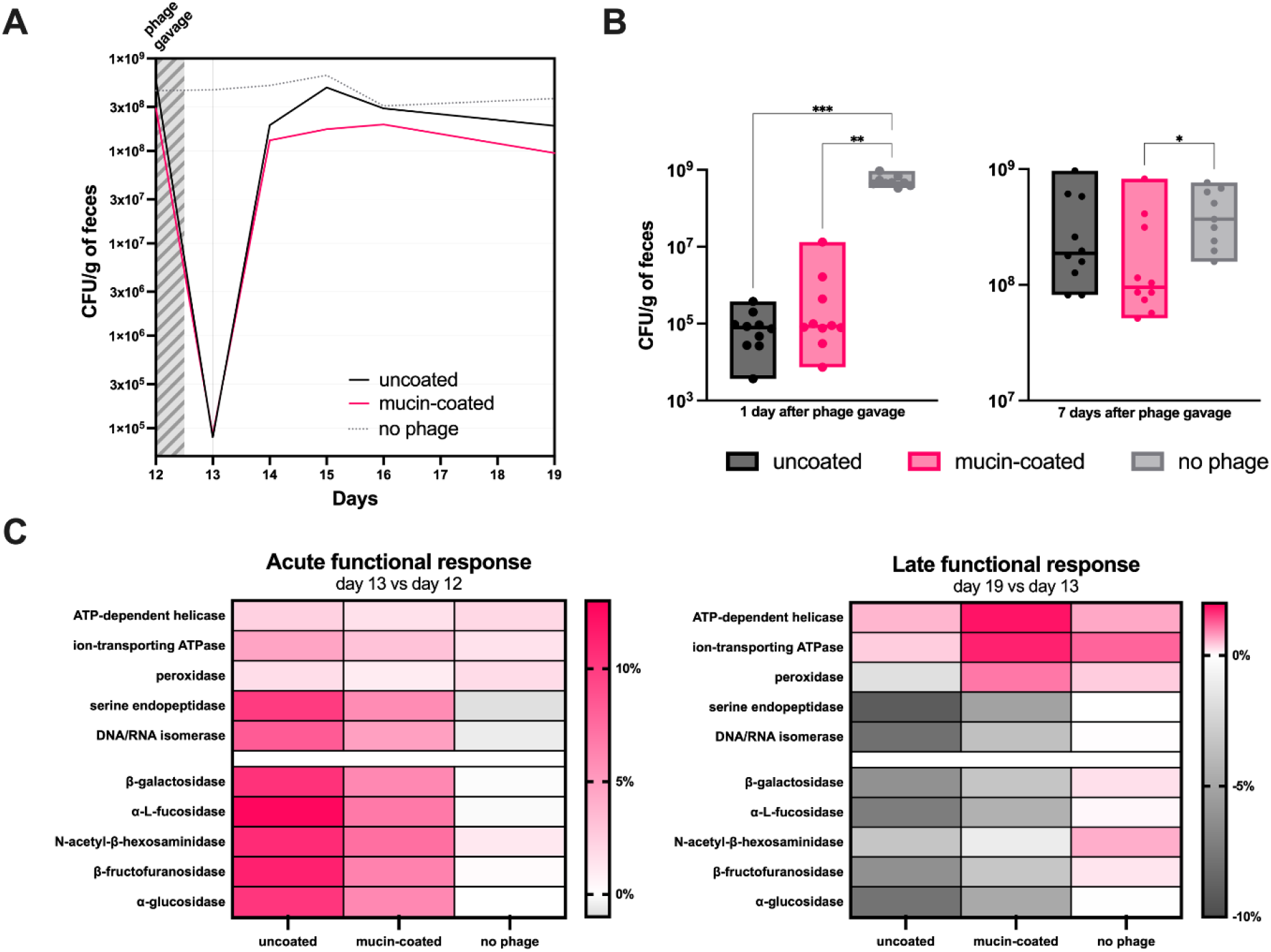
Effects of T4 on *E. coli* colonization and overall predicted microbiome enzyme functions. To assess whether the increased persistence of phages in mice treated with mucin-coated T4 extends to bacterial dynamics, we monitored *E. coli* MG1655 levels and downstream microbiome shifts over time. **(A)** Fecal *E. coli* colonization levels (CFU/g) across the experimental sampling window, displayed as median values for uncoated T4 (*n* = 10), mucin-coated T4 (*n* = 10) and no phage control treatment (*n* = 9). Gray shading indicates the timing of phage gavage. **(B)** Fecal *E. coli* levels (CFU/g) shown for individual mice 1 day after phage gavage (day 13) and 7 days after phage gavage (day 19). Horizontal lines indicate the median with interquartile range. Statistical significance was assessed using Kruskal-Wallis with Dunn’s post-hoc correction. **(C)** Acute and late functional responses of the fecal microbiome following phage treatment. Heat maps show within-group percentage changes in PICRUSt2-predicted enzyme commission (EC) abundances for the three treatment conditions. The acute functional response depicts percent change from day 12 to day 13, and the late functional response change from day 13 to day 19. EC values were first calculated per mouse (*n* = 5 mice per group), then summarized within treatment groups. Color scales are centered at 0% to indicate no change, with positive values (pink) indicating percentage increases and negative values (grey) indicating decreases relative to the preceding timepoint. ECs are grouped by functional category, with stress-associated enzymes shown above mucin-associated glycoside hydrolases.

To assess changes in gut microbiome composition, we performed 16S rRNA gene sequencing on fecal samples collected at the three timepoints mentioned above. For each timepoint and treatment condition, feces from five mice per group were analyzed to ensure balanced representation for downstream community-level analyses. Initial taxonomic profiling showed that the gut microbiome was dominated by members of the *Muribaculaceae* and *Bacteroides* genera across all experimental groups, consistent with their well-established prevalence in SPF mouse microbiota (**Fig. S2**) [24]. Notably, already prior to phage administration, microbiome composition differed between individual mice and across experimental groups within the expected range of baseline heterogeneity for SPF housing conditions (**Fig. S2**). This variability was likely further influenced by variable exposure to streptomycin administered ad libitum via drinking water rather than in a dose-controlled manner, which is known to perturb gut microbiome composition and reduce overall community diversity [25]. As these differences preceded phage treatment, they represent pre-existing biological variable states rather than treatment effects. To focus on treatment-associated dynamics, we quantified microbiome changes as within-group percentage changes over time normalized to corresponding changes in no-phage controls, thereby accounting for background temporal drift.

To identify overall bacterial taxonomic shifts between treatment groups following phage exposure, we performed a SIMPER analysis on relative taxa abundance. This analysis indicated that members of the *Muribaculaceae, Bacteroides, Escherichia-Shigella*, and *Parabacteroides* genera accounted for the largest contributions to between group dissimilarity. In particular, changes in *Escherichia*-*Shigella* abundance reflected the dynamics of the introduced *E. coli* strain following phage treatment, while shifts in *Muribaculaceae* and *Bacteroides* indicated broader alterations among dominant commensal populations in response to phage exposure [24]. To assess treatment-associated taxonomic changes (**Fig. S2**) in more detail, we quantified temporal shifts as within-group percentage changes between consecutive timepoints before treatment vs. day 1 after treatment as acute treatment response (experimental day 12 to day 13), and day 1 after treatment vs day 7 after treatment as late treatment response (experimental day 13 to day 19), respectively, normalized to the corresponding changes observed in no-phage controls to account for background temporal drift. As expected, day 1 after phage treatment, *Escherichia-Shigella* plummeted, indicating substantial *E. coli* killing, with reductions more pronounced in mice treated with uncoated T4 (-16.2%) compared to mucin-coated T4 (-9.2%). This decline was accompanied by expansion of fast-growing taxa, consistent with a rapid occupation of a vacated niche [25]. Specifically, *Bacteroides* increased in both the mice treated with uncoated T4 (+4.1%) and mucin-coated T4 (+6.5%), while *Parabacteroides* showed a pronounced early expansion specifically in the mice receiving uncoated T4 (+26.4%), indicative of a strong opportunistic response. Notably, these taxonomic groups include members capable of degrading mucin-derived glycans, although not all members possess mucin-degrading capacity [26, 27]. At 7 days after phage treatment, *Escherichia-Shigella* showed partial recovery relative to experimental day 13. Increases were more pronounced in mice treated with uncoated T4 (+ 11.3%) than in animals treated with mucin-coated T4 (+5.3%). Concurrently, taxa that had bloomed in the absence of *E. coli* declined again, whereas other mucus-associated taxa expanded: Most notably *Akkermansia*, a specialized mucin-degrading bacterium [26], which increased strongly in the mice treated with mucin-coated T4 (+37.0%) compared to mice treated with uncoated T4 (+14.2%). In contrast, *Alistipes* and unclassified *Muribaculaceae* showed more comparable late phase increases across phage treated groups, the latter representing a taxonomically diverse family with heterogeneous capacities for host-glycan utilization [28].

To investigate functional shifts between treatment groups over time, we inferred predicted enzyme commission (EC) profiles from our 16S rRNA gene sequencing data using PICRUSt2, which estimates total microbial enzymatic potential by linking taxonomic composition to functionally annotated reference genomes. EC abundances were determined per mouse and summarized within treatment groups at each timepoint. Functional changes were quantified as within-group percent changes between timepoints, enabling direct comparison of acute (experimental day 13 vs day 12) and late (experimental day 19 vs day 13) treatment responses (**Fig. 5C**). The ECs exhibiting the strongest treatment-associated variation grouped into two broad functional categories: stress-associated enzymes and enzymes involved in mucin-associated carbohydrate degradation. Cellular stress related enzyme classes included an ATP-dependent helicase (EC 3.6.4.12), ion-transporting ATPase (EC 3.6.3.12), peroxidase (EC 1.11.1.15), serine endopeptidase (EC 3.4.21.89), and DNA/RNA isomerase (EC 5.99.1.2), representing pathways involved in genome maintenance, membrane homeostasis, proteolysis, and oxidative stress responses. Furthermore, these are commonly engaged during lytic phage infection and have been shown to undergo rapid and differential transcriptional regulation in *E. coli* immediately following T4 exposure [29]. Mucin-associated ECs included β-galactosidase (EC 3.2.1.23), α-L-fucosidase (EC 3.2.1.51), N-acetyl-β-hexosaminidase (EC 3.2.1.22), β-fructofuranosidase (EC 3.2.1.52) and α-glucosidase (EC 3.2.1.20), all belonging to glycoside hydrolase families involved in breaking down mucin O-glycans and related carbohydrates [27]. At baseline (experimental day 12), predicted abundances of selected ECs were broadly comparable yet not identical across experimental groups, reflecting baseline heterogeneity in fecal microbiome composition that we also observed at the taxonomic level (**Fig. S2**). Standardized EC z-scores - defined as predicted EC abundances normalized to the mean and standard deviation across samples - are visualized in **Fig. S3** for each timepoint. Because predicted EC profiles are inferred directly from community composition, such baseline functional differences are expected and represent pre-existing ecological variation rather than treatment effects. To account for this heterogeneity and focus on treatment-associated dynamics, we therefore quantified within-group percentage relative changes between timepoints rather than comparing absolute predicted EC abundances across groups.

One day after phage administration (day 13 vs day 12), both phage-treated groups exhibited an acute functional response, characterized by increased predicted abundance of both stress-associated and mucin-associated ECs (**Fig. 5C**). This response was consistently stronger in the mice treated with uncoated T4, with stress-associated ECs increasing by 5-10%, compared to 3-6% in the group treated with mucin-coated T4. Mucin-associated glycoside hydrolases showed a similar pattern, with larger early increases in the mice receiving uncoated T4 (8-13%) than in the mice receiving mucin-coated T4 (5-7%). These acute functional changes coincided with the pronounced (∼4 log_10_) reduction in *E. coli* abundance following phage treatment (**Fig. 5A**), reflecting a substantial ecological perturbation and paralleling early taxonomic niche expansion of fast-growing luminal taxa such as *Bacteroides* and *Parabacteroides* (**Fig. S2**). Accordingly, the observed functional shifts likely reflect indirect consequences of phage mediated reductions in *E. coli* abundance and associated changes in microbial interactions, with phage activity functioning as an upstream driver rather than exerting direct effects on the broader microbiota.

At day 7 (experimental day 19 vs day 13), in the microbiome of the mice receiving uncoated T4, both stress-associated and mucin-associated ECs declined (-6 to -10%), indicating resolution of the acute functional response. In contrast, the microbiome of the mice receiving the mucin-T4 showed continued positive or near-zero changes in stress-associated ECs (0-2%), indicating a sustained functional response rather than resolution. This observation was most pronounced for the ATP-dependent helicase and ion-transporting ATPase, two enzymes with well-established roles in phage-related cellular stress: Helicase linked functions are known to participate in phage-responsive genome integrity surveillance [30], and ATPase-associated ion-homeostasis pathways are characteristic of phage-induced envelope stress, including activation of the phage shock protein response [31]. Mucin-associated glycoside hydrolases followed a similar trend, with abundances showing smaller late-phase decreases in mice treated with mucin-coated T4 compared to the uncoated T4. In contrast, predicted EC abundances in no-phage controls showed minimal changes over time and remained near baseline (0%).

Overall, analysis of predicted EC dynamics reveals a shared acute functional response to phage treatment, followed by distinct late-phase trajectories. Importantly, because predicted EC profiles are derived from taxonomic composition, these functional dynamics reflect underlying microbiome restructuring that is shaped by phage exposure. Uncoated T4 treatment induced a strong but more transient functional perturbation that largely resolved one week after treatment. In contrast, mucin-coated T4 treatment induced a delayed and prolonged functional response, with weaker early changes followed by continued activation of stress-associated functional signatures at later timepoints. This pattern aligns with delayed taxonomic shifts and increased persistence of mucin-coated phages within the gut, consistent with prolonged phage-host interactions.

Together, these data indicate that mucin type II modulates T4-*E. coli* interactions across multiple stages, from delaying initial phage attachment (**Fig. 1**) and protecting virions from environmental stress (**Fig. 3**) to prolonging phage persistence during GI passage (**Fig. 4**). This enhanced persistence of mucin-coated T4 *in vivo* translates into sustained suppression of *E. coli* populations and is accompanied by prolonged and temporally shifted changes in predicted stress-associated functional signatures in the fecal microbiome (**Fig. 5**). Collectively, these findings support a model in which mucin coating extends the duration and ecological impact of T4 phage activity.

## Discussion

Mucosal surfaces constitute a dynamic biochemical environment where phages, bacteria, and host factors interact. This environment is inherently complex, shaped by interdependent physical, chemical, and biological processes, resulting in a tightly interlinked system in which polymer physics, microbial metabolism, and phage behavior modulate one another over time. Our findings place mucin at the center of this system as an active modulator of both phage infection dynamics and phage stability.

The BAM model proposes that phages such as T4 adhere to mucin glycans through capsid-associated domains like Hoc, thereby increasing their retention within mucus and enhancing their likelihood of encountering bacterial hosts [15]. Moreover, the mucin gel architecture physically constrains T4 motion thereby influencing T4-*E. coli* encounter frequency [16]. Our data extend the BAM framework by showing that mucin does not simply enrich phages spatially but also reshapes infection kinetics and downstream ecological consequences over time.

We found that purified mucin type II delays T4 infection, without affecting T4 replication output (**Fig. 1**). Mechanistically, we propose that T4 becomes transiently sequestered within mucin aggregates, consistent with the ability of mucin to form weak, reversible intermolecular clusters through non-covalent mucin-mucin interactions. Such structures form at pH 5.8, closely matching the pH 5.6 of our mucin-supplemented LB [32]. In this state, T4 appears physically restricted, with limited access to host receptors. Importantly, this occlusion is temporary, indicating that mucin-mediated T4 shielding and subsequent release are dynamically coupled rather than static features of the matrix. This is likely caused by gradual rearrangement of weakly cross-linked mucin networks, consistent with the reversible nature of mucin-mucin and mucin-phage interactions [13, 15]. At the same time, active bacterial processing of mucin represents a plausible complementary mechanism, particularly over longer time, in more complex microbial environments such as the gut microbiome which is rich in mucin degraders [27]. While our data does support a model in which *E. coli* MG1655 actively remodels or degrades the mucin coating, progressively releasing occluded virions and enabling infection, most *E. coli* lineages cannot degrade native intestinal mucin, due to its heavily glycosylated O-glycan chains that require specific glycosidases that *E. coli* generally lacks [33]. An exception is entero-toxigenic *E. coli*, ETEC, producing mucin-targeting proteases such as EatA that directly cleave native mucin [34]. In contrast, we used purified mucin type II, which is substantially less glycosylated and structurally simpler. Despite apparently lacking specialized enzymes for degradation of native mucin, we found that *E. coli* MG1655 can use mucin type II as sole carbon source (**Fig. 2**), in line with a previous report [33].

Overall, our findings support a model in which mucin shields T4 phages, reducing access to bacterial receptors and delaying overall infection. Essentially, the mucin provides environmental protection and provides a slow-release mechanism. Our proposed mechanism extends prior *in vitro* observations showing that mucin can modulate short-term T4-*E. coli* encounter dynamics by altering viscosity and spatial organization [35]. It was reported that increased agitation and phage concentration partially restored T4 activity, consistent with a physical limitation on encounter rates rather than intrinsic bacterial resistance. Further, studies have proposed that spatiotemporal constraints on phage-host encounters could, in some systems, be alleviated through processing of matrix components by phage-encoded carbohydrate-active enzymes. This hypothesis has been raised in the context of phage-dependent mucin growth phenotypes observed in *Y. enterocolitica* [20]. Phages encode depolymerases, lysins, and related glycan-active enzymes capable of modifying complex carbohydrates produced by bacteria [36]. Whether phages directly modify mucin substrates remains an open question, as this has not been experimentally shown so far.

Similar principles apply at mucosal surfaces, where mucus and biofilm form an integrated polymeric environment that shapes phage-host encounters. Bacterial biofilms restrict phage mobility and receptor access through their extracellular polymer matrix, and some phages have evolved the ability to degrade the matrix gaining access to the bacteria [37]. The mucofilm model extends this analogy to mucosal systems by treating host-derived mucus and bacterial biofilm as a single, integrated polymeric environment [38]. Within this environment, the combined architecture and dynamics of mucus and biofilm jointly shape phage-bacterium interactions, despite imposing constraints distinct from those of classical biofilms.

While mucin-mediated occlusion delays T4 adsorption, we also found that mucin coating actively shields virions and allows them to persist under acidic and proteolytic stress (**Fig. 3**). This mechanism parallels phage encapsulation strategies that prolong phage survival through physical protection. Such approaches include chitosan nanoparticles [39], liposomes [40, 41], and alginate hydrogels [42] which all improve phage stability or enable targeted phage release. A particularly relevant example comes from a recent study that encapsulated *Salmonella* phages in hydrogel microspheres [43]. In that work, free phages were rapidly inactivated by a pH 2.5 gastric challenge, whereas hydrogel-encapsulated phages remained infectious, demonstrating substantial acid protection.

Beyond increased stability of virions *in vitro*, we observed enhanced *in vivo* persistence of mucin-coated T4 phages (**Fig. 4**) that correlated with prolonged suppression of *E. coli* levels (**Fig. 5**). This observation is similar to the study in which encapsulated *Salmonella* phages were found to persist considerably longer: while free phages vanished from the murine gut within 8 h due to intestinal excretion and degradation, encapsulated phages remained detectable at 8-12 h before clearance by 24 h [43]. In parallel, recent work has shown that mucin-associated phages exhibit increased recovery, with approximately 10-fold higher phage titers after 24 h in mucin-containing media than in LB and sustained phage levels in fecal samples up to 24 h, whereas blocking phage-mucin interactions via Hoc antibody treatment resulted in loss of detectable phage in feces within 12 h, consistent with enhanced mucus-associated retention and antimicrobial effects [21].

Although mucin coating delayed infection initiation *in vitro*, overall maximal phage replication capacity was indistinguishable between uncoated- and mucin-coated phages (**Fig. 1**), indicating that mucin does not impair phage amplification once infection is established. At the same time, mucin-mediated T4 protection against acidic and proteolytic stress *in vitro* (**Fig. 3**) likely contributes to, but does not fully explain, the enhanced T4 recovery after GI passage (**Fig. 4**). The absence of detectable T4 sensitivity in *E. coli* isolates recovered 1 day after phage gavage (**Fig. 4D**), indicates rapid emergence of phage resistance *in vivo*. This is expected, as single-phage treatments impose a high selection pressure, selecting for resistant bacterial subpopulations within hours in animal models [44]. However, resistance was assessed only in fecal *E. coli* isolates challenged against the ancestral laboratory T4 phage, capturing bacteria that had transited the gut rather than the full intestinal population. As such, this assay does not resolve the ongoing co-evolutionary dynamics between phage and bacteria *in vivo*, including potential diversification of phage populations or spatially structured *E. coli* resistance, e.g. by physical shielding within mucofilm, in the gut. Such T4-susceptible subpopulations of *E. coli* likely contribute to the prolonged phage recovery, in combination with T4 adherence to- and continuous release from the mucus surface.

Importantly, additional biological factors may influence phage abundance in the gut, including phage receptor regulation. Although we did not detect mucin-induced changes in the abundance of the T4 receptor OmpC *in vitro* (**Fig. 1C**), differential receptor regulation in the complex murine intestinal environment cannot be ruled out. In *E. coli, ompC* is transcriptionally upregulated under high osmolarity and in intestinal environments, resulting in increased OmpC levels [45, 46]. Thus, we cannot exclude the possibility that the mucin-mediated increased T4 abundance in murine pellets reflects a combination of enhanced T4 stability, increased retention, and context-dependent replication, representing a limitation to our present study. Entirely disentangling the relative contributions of mucin-mediated stability, retention, and context-dependent replication will require future studies capable of resolving these processes independently in the intestinal environment.

Further, at the microbiome level, we detected prolonged enrichment of predicted stress-response and mucin-utilization enzyme functions in the mice receiving mucin-coated T4, whereas these functional signatures largely resolved in control T4-treated mice (**Fig. 5C**). Because these profiles were inferred from 16S rRNA gene data using PICRUSt2, they primarily reflect underlying taxonomic shifts rather than enzyme enrichment within individual taxa. Notably, the acute enrichment of stress-response and mucin-degradation functions following phage treatment coincided with pronounced restructuring of the microbial community at the early post-treatment timepoint (day 13). The T4-driven *E. coli* loss vacated a niche, relieving competitive pressure and freeing both physical space and resources. Nonetheless, the persistence of these predicted functions in the treatment group receiving mucin-coated T4 indicates a prolonged, mucin-enhanced effect of T4-induced functional remodeling at the microbiome level.

Beyond effects on phage persistence, mucin-rich environments reshape phage-bacterial-host interactions by altering bacterial behavior itself. Mucin activates *F. columnare* virulence and increases initial phage susceptibility, but also promotes phage defense mechanisms such as CRISPR adaptation, demonstrating a dynamic balance between phage susceptibility and resistance [17, 18]. In *S. mutans*, mucin similarly increases phage susceptibility and underscores the role of the mucosal environment in shaping bacterial growth, phage sensitivity, and resistance phenotypes [19]. Mucin exposure also influences phage resistance evolution in *Y. enterocolitica*, with resistance arising through mutations linked to virulence, quorum sensing, and antibiotic resistance, pointing to potential fitness trade-offs during adaptation under phage pressure [20]. Together, these studies highlight that mucosal effects on phage-host interactions are highly context dependent, underscoring the need for system-specific characterization of phage-bacterial-mucin interactions.

Although our study uses purified mucin type II, which lacks the full structural complexity of native mucus, it demonstrates that mucin glycopolymer environments impose spatiotemporal constraints on phage infection dynamics. Mucin provided a stabilizing microenvironment that protected T4 from acidity and proteolysis *in vitro*, enhanced persistence in the murine gut and increased fecal phage recovery. Together, our data suggest that mucin shapes T4 phage biology by delaying infection initiation while extending virion stability and persistence. While the use of mucin as adjunct to oral phage delivery has been proposed previously [47], our data provide direct mechanistic and *in vivo* support for these efforts, highlighting how mucin-based or hybrid biological-synthetic matrices may be leveraged to extend phage persistence, after a single oral dose, and tune infection dynamics in GI environments.

## Acknowledgements

We thank Joana Paula Gomes Amaro for assistance with fecal sample preparation and PFU/CFU assays, and Eva Oros for assistance with phage resistance assays. Further, we acknowledge the Core Facility for Integrated Bioimaging, Faculty of Health and Medical Sciences, University of Copenhagen for TEM imaging. K.C.K. was supported by the EMBO Scientific Exchange Grant 11375 and N.M.H.-K. was supported by the Independent Research Fund Denmark grant 1054-00099B.

## Author contributions

**KCK** performed all main laboratory experiments, developed methodology, analyzed data, and wrote the original manuscript. **AR** performed initial *in vitro* work that guided the final experimental design and phase contrast microscopy. **RBG** led *in vivo* data collection. **MM** provided microscopy resources and supervised imaging work. **KBX** supervised and housed all *in vivo* work. **JJM** contributed to early conceptual framing and supervision. **NMHK** served as lead PI, overseeing and supporting all aspects of the project. All authors contributed to the final manuscript and approved the final version.

### Materials and Methods Bacterial Culturing

*E. coli* ARO43, a streptomycin-resistant (strep^R^) MG1655 strain [25] (hereafter *E. coli* MG1655) was routinely grown at 37 °C with 300 rpm shaking in LB medium (10 g/L tryptone, 5 g/L yeast extract, and 10 g/L NaCl). When appropriate, overnight cultures were diluted in fresh LB to a specified optical density (OD_600_).

### Plaque Assays

Phage titers were determined using the double-layer agar or lawn overlay method. Soft agar (LB with 0.75% agar) containing a mid-log *E. coli* MG1655 culture was poured onto LB agar plates to form a bacterial lawn. Phage samples were either mixed with the soft agar before pouring or spotted onto the solidified lawn, depending on the assay. Plates were incubated at 37 °C overnight, and plaques were quantified the following day.

### Mucin Coating of Phages

For phage coating, T4 phage stocks were diluted to an appropriate concentration (typically 10^7^ PFU/mL) in LB medium or LB supplemented with 1% (w/v) porcine gastric mucin type II (Sigma Aldrich). Phage suspensions were incubated at 37 °C with constant shaking at 300 rpm for 30 min to allow interaction between phages and mucin polymers. This incubation step was performed prior to bacterial exposure to ensure that phages were equilibrated with mucin.

### Adsorption Assays

Phage suspensions were diluted to a final concentration of 10^5^ PFU/mL in either LB medium or LB medium supplemented with 1% (w/v) mucin. To initiate adsorption, 100 µL of OD_600_ of 0.1 *E. coli* MG1655 culture was added to 1 mL of the phage suspension and briefly vortexed. Samples were incubated at 37 °C without agitation. At defined time points, aliquots were treated with 0.1 volumes of chloroform. Free phages were quantified by plaque assays.

### Western Blot

*E. coli* MG1655 was grown overnight in LB medium and subsequently back-diluted into either LB or LB supplemented with 1% (w/v) mucin then grown and harvested simultaneously at the indicated optical densities. Cell pellets were resuspended in Laemmli SDS sample buffer and denatured at 70 °C for 10 min. Samples were separated by SDS-PAGE on a 4-15% Mini-PROTEAN TGX stain-free polyacrylamide gel (Bio-Rad) and blotted onto an Amersham Protran nitrocellulose 0.45 µm membrane. After blocking in Tris-buffered saline with Tween 20 (TBST) and 5% skimmed milk, the membrane was cut and incubated with either the OmpC polyclonal HRP-conjugated antibody (catalogue no. A64455-100; 1:3,000 dilution in TBST with 5% skimmed milk) or the monoclonal Anti-RpoB antibody (catalogue no. ab191598; Abcam; 1:3,000 dilution in TBST with 5% skimmed milk) for 1 h. After three washes with TBST, the Anti-rabbit IgG HRP conjugate antibody (catalogue no. W4011; 1:10,000 dilution in TBST with 5% skimmed milk) was used as the secondary antibody for detection of the Anti-RpoB antibody. The membranes were washed in TBST and developed using SuperSignal West Femto maximum sensitivity substrate (catalogue no. 34095; Thermo Scientific). The relative band intensities were quantified in Image Studio.

### Carbon Utilization Assay

Overnight cultures were back-diluted to an OD600 of 0.01 and grown for 1.5 h. Cells were washed three times and subsequently resuspended in M9 supplemented with 2mM MgSO_4_ and 0.1mM CaCl_2_ without, or with, 0.4% (w/v) glucose, 1% (w/v) mucin, or resuspended in LB. The cultures were incubated at 37 °C for 20 h with intermittent shaking in a plate reader (Synergy H1 BioTek Instruments Inc.). Media-only controls for each condition were included for background subtraction.

### Phase-Contrast Microscopy and Time-Lapse Imaging

*E. coli* MG1655 was grown overnight in LB, diluted 1:20 in M9 minimal medium supplemented with 0.4% (w/v) glucose, and inoculated into M9 supplemented with 1% (w/v) mucin. Cells were transferred to *µ*-Slide I Luer 3D (ibidi) for imaging. Time-lapse phase contrast microscopy was performed at room temperature using an Olympus Corporation BX61 microscope equipped with an Olympus DP74 camera at a 40x phase contrast objective. Images were acquired every 2 min for 10 h.

### Transmission Electron Microscopy

T4 phages were treated with 1% (w/v) mucin in LB medium prior to imaging. Samples were applied to glow discharged carbon coated copper grids using double sample application with 1 min incubation time to increase material retention on the grid. Excess liquid was gently blotted, and grids were negatively stained with 2% (w/v) uranyl acetate. Imaging was performed using a FEI Morgagni 268D transmission electron microscope at a magnification of 180,000×.

### Stability Assays

After coating the phages at a concentration of 10^7^ PFU/mL, suspensions were diluted in buffer (0.9% NaCl, 20 mM CaCl_2_, 50 mM Tris-HCl) to a final concentration of 10^5^ PFU/mL. For pH-dependent stability assays, buffer aliquots were adjusted to pH 7.0 or pH 3.0, and samples were incubated at 37 °C with shaking at 300 rpm for up to 60 min. For α-chymotrypsin assays, the buffer was adjusted to pH 8.0 and supplemented with 3 mg/mL α-chymotrypsin where indicated. Samples were incubated at 37 °C for up to 48 h without agitation. Phage viability was quantified using plaque assays. The protocol was adapted from Majewska *et al*. [23].

### Ethical Statement

All experiments involving mice were approved by the Institutional Ethics Committee at the Gulbenkian Institute for Molecular Medicine (GIMM) and by the Portuguese National Entity that regulates the use of laboratory animals (DGAV - Direção Geral de Alimentação e Veterinária (license reference: 015190). All experiments conducted on animals followed the Portuguese (Decreto-Lei n° 113/2013) and European (Directive 2010/63/EU) legislations, concerning housing, husbandry, and animal welfare.

### Mouse Colonization and Phage Treatment

All mice were 6-to 8-week-old male C57BL/6J conventionally raised under SPF conditions and supplied by the Rodent Facility at GIMM. Animals had ad libitum access to food (Rat and Mouse No. 3 Breeding diet; Special Diets Services, product no. 801030) and water. Mice were maintained at 20-24 °C and 40-60% humidity on a 12 h light-dark cycle. Sample size was chosen in accordance with institutional guidelines and principles for the humane use of animals in research. The experiment was performed twice following the same protocol. At the start of each experiment, mice were randomly assigned to experimental or control groups and individually housed in isocages. From this point, mice received streptomycin (5 g/L) in drinking water ad libitum, replaced every 2-3 days, to allow and maintain intestinal *E. coli* colonization. All mice were orally gavaged with 100 µL 1x PBS containing 10^8^CFU of *E. coli* MG1655 strep^R^. After confirming stable colonization over 10 days based on fecal CFU monitoring, mice were orally gavaged with 10^7^ PFU uncoated phages in 100 µL suspension (*n* = 10), 10^7^ PFU mucin-coated phages in 100 µL suspension (1 mg total mucin per mouse, *n* = 10), or 100 µL sterile 1x PBS as a solvent control (*n* = 9). Fresh fecal samples were collected for enumeration of *E. coli* MG1655 CFU and T4 PFU, and for DNA extraction. Fecal samples were collected over 27 days. Fecal samples for DNA analyses were weighed and stored at - 80 °C until further processing.

### Bacterial and Phage Enumeration from Feces

For CFU enumeration, fecal pellets were weighed, mechanically disrupted with a motorized pellet pestle in 1 mL sterile 1x PBS, and serially diluted in 1x PBS, spotted onto LB and 100 µg/mL strep LB plates. For PFU enumeration, fecal suspensions were treated with 0.1 volumes of chloroform to lyse bacterial cells and then serially diluted in 1x PBS. Phages were quantified by plaque assays. CFU and PFU plates were incubated overnight at 37 °C and counted the following day.

### Phage resistance assays

Fecal pellets were stored at -80 °C in 1x PBS supplemented with 20% glycerol and streaked onto 100 µg/mL LB-strep plates. Material from the resulting streaked growth was used to inoculate overnight cultures in 100 µg/mL strep-supplemented LB to maintain selection for *E. coli* MG1655. Overnight cultures were diluted in LB-strep to an initial OD_600_ of 0.01 (∼10^7^ CFU/mL). T4 was added to a final concentration of ∼2 × 10^5^ PFU/mL, corresponding to a multiplicity of infection of 0.02. Cultures with or without phage were dispensed into 96-well plates, and bacterial growth was monitored over time using a plate reader by measuring OD_600_ to assess susceptibility to phage infection.

### DNA Extraction, 16S rRNA Sequencing and Fecal Microbiota Analysis

DNA was extracted from fecal pellets using the QIAamp Fast DNA Stool Mini Kit (Qiagen) according to the manufacturer’s instructions, with mechanical disruption performed using a motorized pestle, and eluted in 50-100 µL ATE buffer. DNA concentration and purity were assessed using a Qubit 2.0 Fluorometer and a NanoDrop 1000 spectrophotometer. Amplification of the V4 region of the bacterial 16S rRNA gene, library preparation, sequencing, and primary bioinformatic processing were performed by Novogene. Libraries were sequenced on an Illumina NovaSeq platform to generate 250 bp paired end reads. Sequence data were processed by Novogene using a QIIME2 based pipeline, including quality filtering, read merging, denoising and amplicon sequence variant inference with DADA2, and taxonomic assignment using the SILVA database.

## Supplementary Files

**Figure S1.**
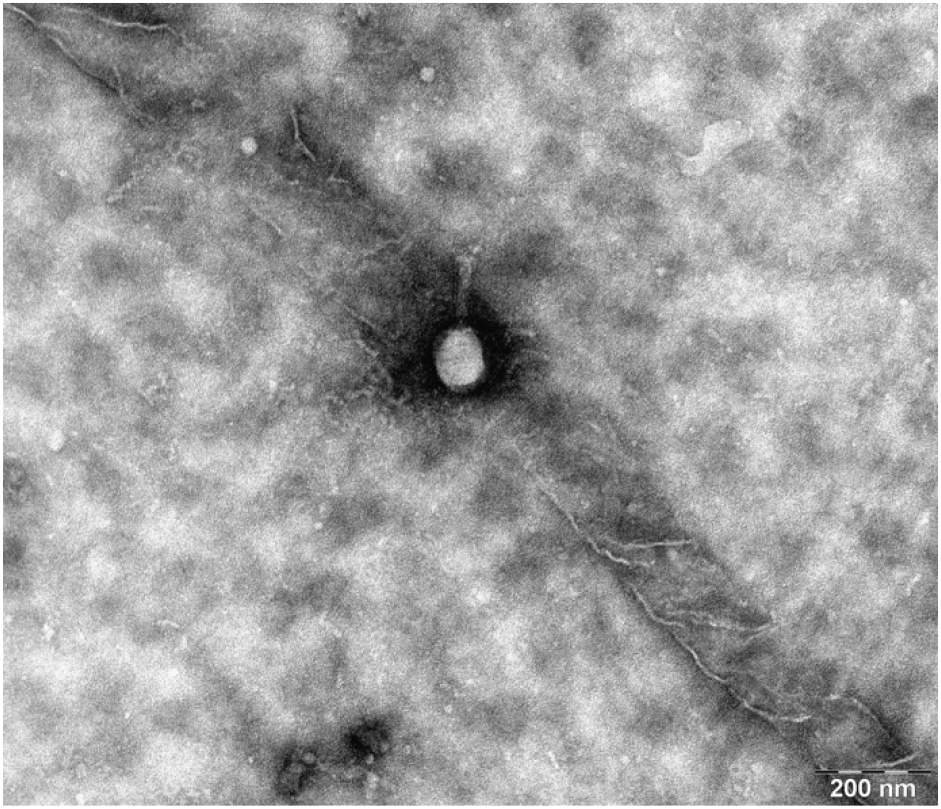
TEM image of mucin-coated T4. T4 phages were precoated with 1% mucin. A T4 phage is associated with a mucin filament. Electron-dense, amorphous material is observed surrounding the phage capsid, consistent with collapsed mucin aggregates, whereas the tail structure appears largely unobstructed. Scale bar 200 nm.

**Figure S2.**
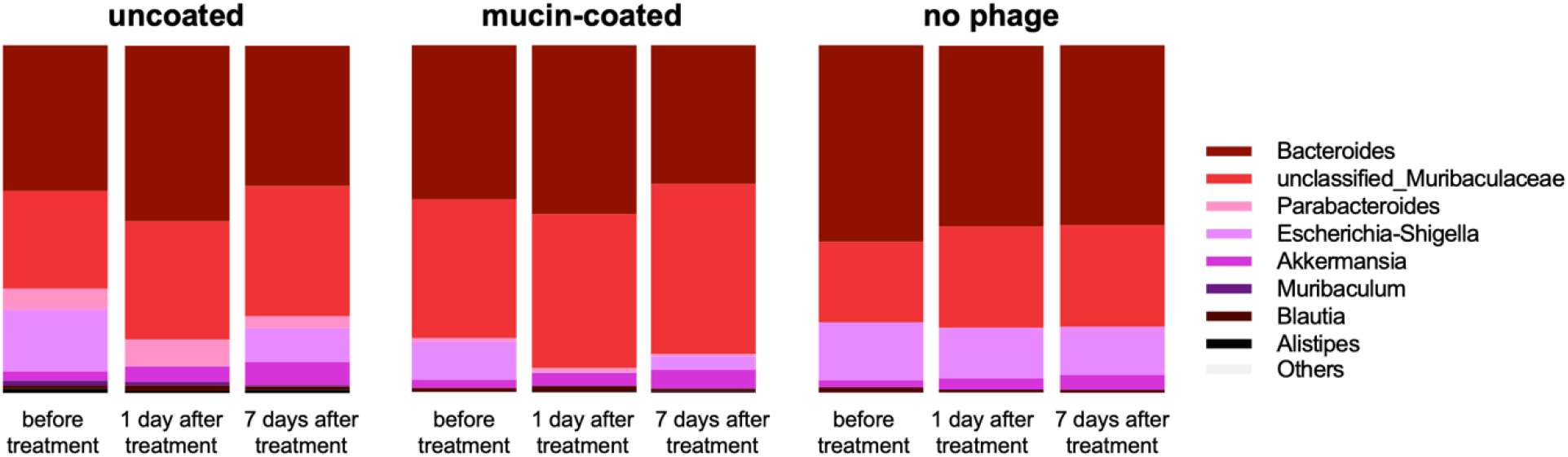
The effect of phage-treatment on fecal microbiome taxonomic shifts. To assess whether differences in T4 formulation influence downstream microbiome dynamics following *E. coli* attack, we monitored changes in the relative abundance of dominant bacterial taxa over time. Stacked bar plots depict mean genus level relative abundances derived from 16S rRNA gene sequencing of fecal samples collected before phage treatment (day 12), 1 day after phage gavage (day 13), and 7 days after phage gavage (day 19) from mice treated with uncoated T4 phage, mucin-coated T4 phage, or no phage as control. Taxa shown include *Bacteroides*, unclassified *Muribaculaceae, Parabacteroides, Escherichia*-*Shigella, Akkermansia, Muribaculum, Blautia, Alistipes*, and remaining low abundance taxa grouped as Others.

**Figure S3.**
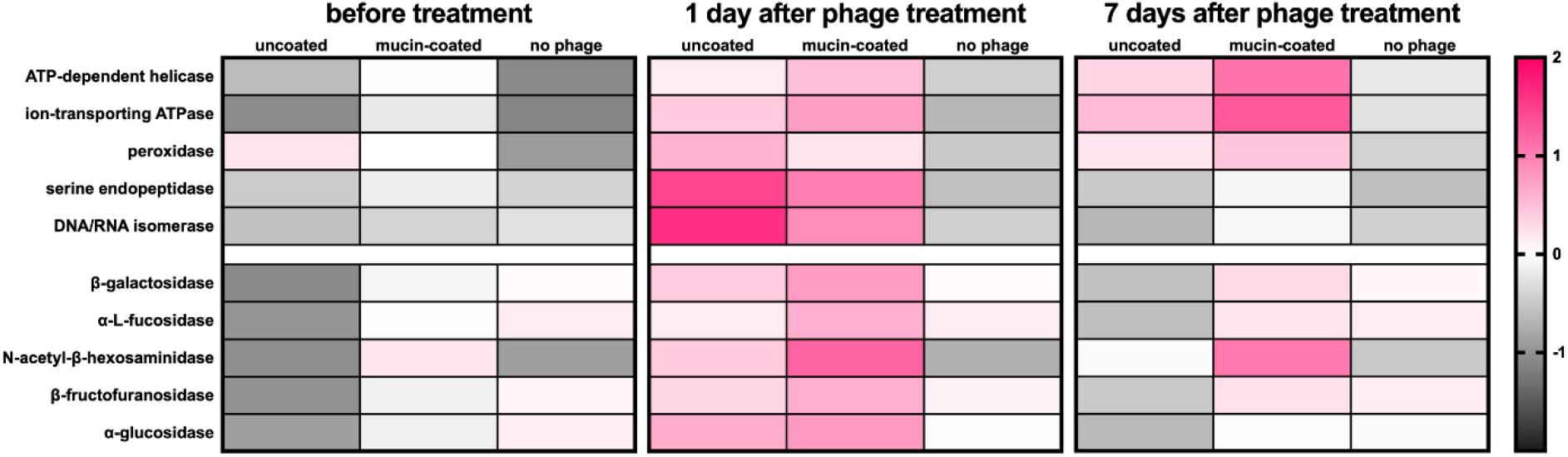
Overall predicted microbiome-wide enzymatic profiles. Heatmaps of PICRUSt2-predicted EC activities in uncoated T4, mucin-coated T4, and no-phage treatment groups across three timepoints: 1 day before phage treatment (day 12), 1 day after phage gavage (day 13), and 7 days after phage gavage (day 19). Values represent standardized z-scores of predicted EC abundances, calculated by centering and scaling across all samples, and are shown to provide context for baseline functional states and absolute shifts over time. Data represent group means pooled from *n* = 5 mice per condition. Color scale indicates deviation from the cross-sample mean, ranging from -2 (lower relative abundance, grey) to +2 (higher relative abundance, pink). This figure complements the percentage change analyses presented in **Fig. 5C**.

